## Supplemental figures for "Cell type determination for cardiac differentiation occurs soon after seeding of human induced pluripotent stem cells"

### Supplementary figure 1

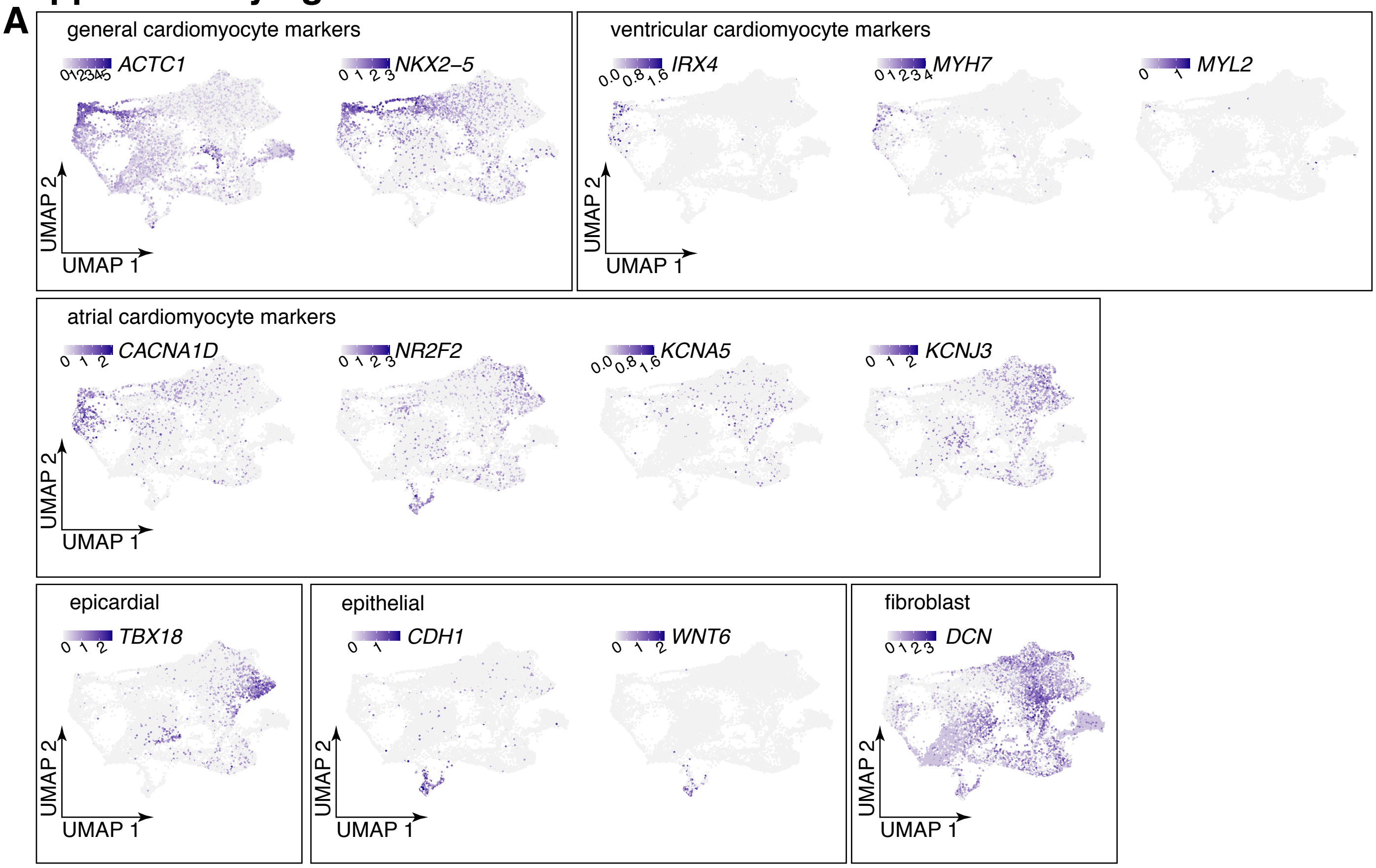


**Supplementary figure 1. Localization of additional cell type markers following cardiac directed differentiation in day 14 differentiated cells.** (A) Maintaining the organization provided by UMAP, we recolored each cell by its expression of a number of canonical cell type markers; some of these are also listed in the heatmap in Figure 1C while others are important in cardiac biology.

### Supplementary figure 2

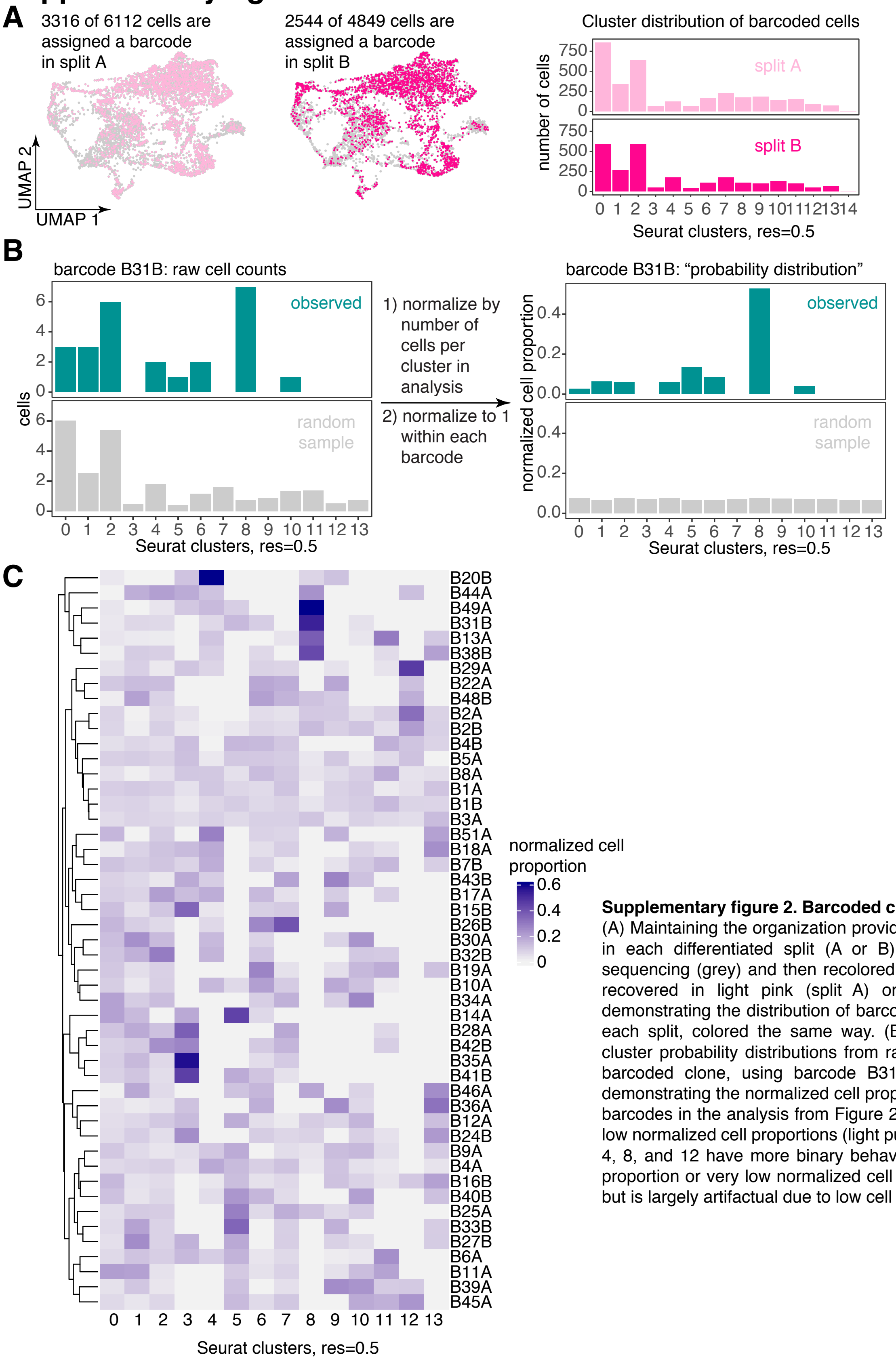


**Supplementary figure 2. Barcoded clone distribution across clusters.** (A) Maintaining the organization provided by UMAP, we recolored all cells in each differentiated split (A or B) profiled by 10X single-cell RNA sequencing (grey) and then recolored the cells for which a barcode was recovered in light pink (split A) or magenta (split B). Bar graphs demonstrating the distribution of barcoded cells across Seurat clusters in each split, colored the same way. (B) Explanation of how we derived cluster probability distributions from raw cell cluster distribution for each barcoded clone, using barcode B31B as an example. (C) Heatmap demonstrating the normalized cell proportion across Seurat clusters for all barcodes in the analysis from Figure 2. Most clusters are associated with low normalized cell proportions (light purple) across barcodes, but clusters 4, 8, and 12 have more binary behavior (i.e. either high normalized cell proportion or very low normalized cell proportion). Cluster 3 does as well but is largely artifactual due to low cell numbers in that cluster.

### Supplementary figure 3

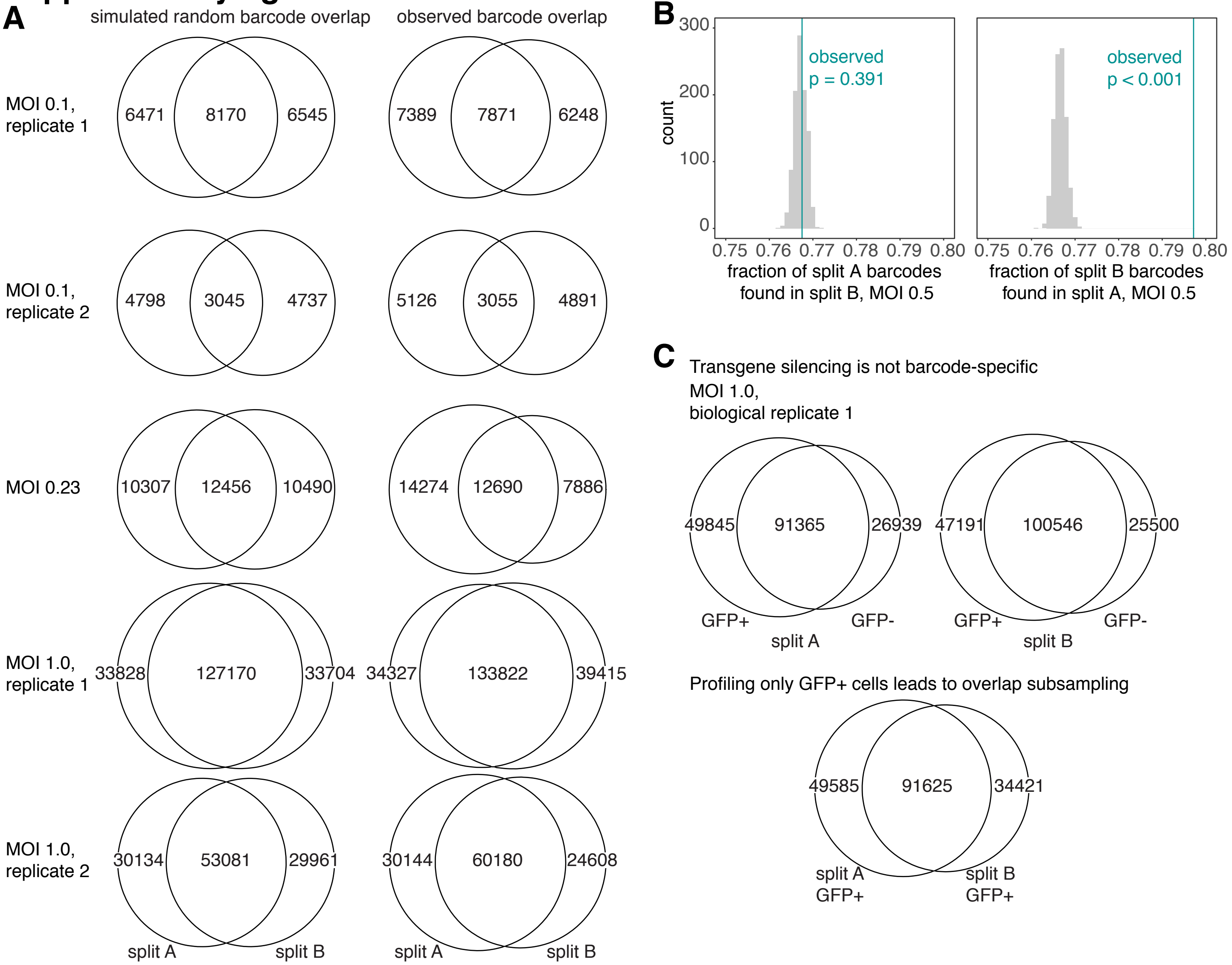


**D**

| Barcode name | Cells in differentiation A | Cells in differentiation B |
|---|---|---|
| B1 | 197 | 301 |
| B2 | 155 | 159 |
| B3 | 186 | 0 |
| B4 | 92 | 27 |
| B5 | 96 | 7 |
| B6 | 83 | 0 |
| B7 | 3 | 73 |
| B8 | 59 | 10 |
| B9 | 68 | 0 |
| B10 | 57 | 5 |
| B11 | 48 | 0 |
| B12 | 48 | 0 |
| B13 | 47 | 0 |
| B14 | 24 | 17 |
| B15 | 0 | 39 |
| B16 | 0 | 39 |
| B17 | 38 | 0 |
| B18 | 23 | 12 |
| B19 | 34 | 0 |
| B20 | 2 | 31 |

**Supplementary figure 3. Neither survival nor transgene silencing are barcode-specific.** (A) Comparison of the simulated random barcode overlap across splits with the observed barcode overlap from the splits differentiated from hiPS cells from all other transduction conditions (MOI 0.1, MOI 0.23, MOI 1.0). (B) Statistical analysis associated with the fraction of overlapping barcodes found after differentiation of cells transduced with an MOI of ~0.5 (Figure 3B). We found that there was no significant difference in overlap for split A. While we did find a significant difference in overlap for split B, the effect size of this difference was such that it may have stemmed from technical variation in barcode recovery and was deemed unlikely to be biologically relevant. (C) Venn diagrams demonstrating the overlap in barcodes found in sorted GFP positive and GFP negative subpopulations following differentiation for each split from cells initially transduced at an MOI of ~1.0. Profound overlap between GFP positive and GFP negative subpopulations suggests that GFP transgene silencing does not occur in an integration-specific or clone-specific manner. (D) Top 20 barcode clones comprising the most total cells across both splits, noting how many were recovered from each split. Pink (light pink for split A, magenta for split B) backgrounds associated with the barcodes that are represented on UMAP in Figure 3C.

### Supplementary figure 4

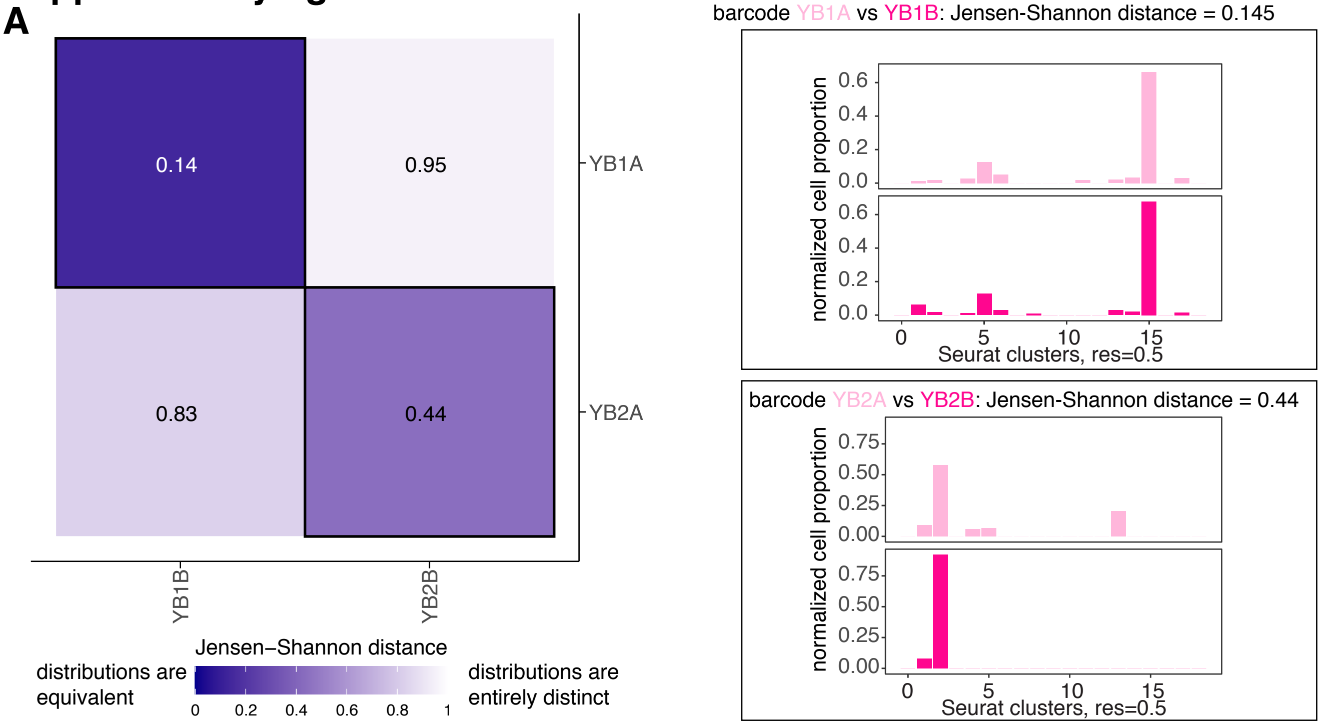


**Supplementary figure 4. Jensen-Shannon distance captures known heritable predetermination of expression state.** (A) Heatmap of pairwise Jensen-Shannon distances between cells associated with 2 barcodes in split A and the cells associated with the same 2 barcodes in split B from a dataset previously shown to have heritable predetermination of the final cell state (vemurafenib-treated melanoma cells). Separated clones sharing a barcode (bolded outline along the diagonal) had much smaller Jensen-Shannon distances than clones labeled by distinct barcodes (off the diagonal), demonstrating their similarity. This similarity is also visible in the comparison for each barcode of bar graphs for observed cluster probability distribution in split A (light pink) and the observed cluster probability distribution in split B (magenta). Cluster probability distributions are visually similar between separated clones and visually distinct across cells labeled by different barcodes.
